# Maize residues changes soil fungal composition and decrease soil microbial co-ocurrence networks complexity

**DOI:** 10.1101/703967

**Authors:** José F. Cobo-Díaz, Fabienne Legrand, Gaétan Le Floch, Adeline Picot

**Author notes:** Corresponding author: José F. Cobo-Díaz.

## Abstract

*Fusarium graminearum* (*Fg*) can cause different diseases in cereals and maize crops worldwide, and a correct management of previous crop residues could decrease disease incidence and/or severity. Bacterial, fungal and *Fusarium* communities were studied by metabarcoding approach in 8 agricultural fields with wheat-maize rotation system in Brittany, France, during three years. Additionally, shift in microbial communities were evaluated under mesocosm experiments in soils amended or not with maize residues and/or *Fg* isolate. Bacterial communities composition were highly influenced by crop soil origin in both environmental and mesocosm soils, while bacteria co-occurrence network complexity was decreased by maize residues in environmental samples and *Fg* treatment in mesocosm samples. Maize residues altered slightly bacteria-fungi co-occurrence networks, while all treatments on mesoscosm experiments showed lower complexity in bacteria-fungi networks than Control Soil treatment. A clear input of fungal genera *Epicoccum*, *Fusarium*, *Vishniacozyma*, *Articulospora*, *Papiliotrema*, *Sarocladium*, *Xenobotryosphaeria*, *Ramularia*, *Cladosporium*, *Cryptococcus* and *Bullera* from maize residues to soil were observed for both environmental and mesocosm samples. Moreover, an increase of *F. graminearum* and *F. avenaceum* was observed in soils whe maize residues were presented. Finally, microbial co-occurrence networks reported some OTUs significant correlated to *Fusarium* spp. OTUs, such as those assigned to *Epicoccum*, *Vishniacozyma* and *Sarocladium* fungal genera, previously reported as efficient biocontrol agents versus *Fusarium* spp. Moreover, a decrease of complexity was observed for soil bacterial and bacterial-fungal networks due to maize addition in both environmental and mesocoms communities.

## INTRODUCTION

*Fusarium* Head Blight (FHB) is a devastating fungal disease of small-grain cereals, including wheat, caused mainly by members of *Fusarium* complexes (Bateman *et al*. 2007; Dean *et al*. 2012). Beyond significantly reduced yields, the main consequence is grain contamination with mycotoxins, including type A and B trichothecenes, produced by toxigenic *Fusarium* spp. (Smith *et al*. 2016; Tralamazza *et al*. 2016). Among *Fusarium* spp. responsible for FHB, *F. graminearum* (*Fg*) is considered the fourth most economically-important plant fungal pathogen (Dean *et al*. 2012), and is within the most frequent species associated to FHB in Europe, besides *F. culmorum, F. avenaceum* and *F. poae* (Xu *et al*. 2005; Hellin *et al*. 2016). Moreover, a shift from *F. culmorum* to *F. graminearum* as the main *Fusarium* species has been recently observed in European cereal crops (Nielsen *et al*. 2011; Scauflaire *et al*. 2011). Climatic change is the principal hypothesis put forward, although the increase in maize-wheat rotation crops may also contribute to this increase in *F. graminearum* at the expense of *F. culmorum* (Nielsen et al, 2011). FHB control represents a major scientific challenge given the multiplicity of causal agents and the complex mechanisms leading to mycotoxin contamination.

The life cycle of *Fusarium* spp. and especially *F. graminearum* is relatively well known (Champeil *et al*., 2004). *F. graminearum* is able to survive for several years saprophytically in soil and especially on crop residues which provide a carrier and nutrients necessary for its growth (Leplat *et al*., 2013). Based on mesocosm experiments, the latter authors demonstrated a higher Fg growth on soil amended with residues than on bare soil while, among crop residues, higher growth was found on soil amended with maize residues, which provided the best carrying capacity over wheat or rapeseed residues (Leplat *et al*., 2016). Similar observations were obtained from field studies (Schaafsma *et al*., 2005; Blandino *et al*., 2010). Overall, the risk of FHB is recognized to be higher when crop residues are left on the soil surface and with direct sowing (Maiorano *et al*., 2008). Therefore, residues, and soil after residue decomposition, are considered as the primary source of inoculum responsible for FHB. In spite of this, the composition and diversity of *Fusarium* spp. in these components have received much less attention than in grains.

Although higher incidence of FHB events were found in maize-wheat cropping systems, especially under minimum tillage practices (Dill-Macky and Jones 2000; Schöneberg *et al*. 2016; Cromey *et al*., 2002; Edwards & Jennings, 2018; Vogelgsang *et al*. 2011), there is some evidences of *Fusarium* spp. survival reduction due to the effect of maize residues microbiota. For instance, Bateman *et al*. (2007) found that chopped maize tops appeared to suppress stem-base disease in wheats, while increasing the presence of *F. graminearun* in the crop, suggesting some microbial activity for disease suppression was occurring in maize residues. This was supported by previous finding of efficient BCAs, in both in vitro and greenhouse trials, isolated from maize tissues (Mousa *et al*. 2015), root rhizosphere (Abiala *et al*. 2015) and residues (Luongo *et al*., 2005; Singh *et al*., 2009).

By colonizing previous crop residues and soil, *Fusarium* spp. can therefore interact with the microbiota associated with these components. Recent advances in next generation sequencing technologies (NGS) has allowed researchers to deepen our knowledge of bacterial and fungal communities in both soils (Chen *et al*. 2015; Zhao *et al*. 2016) and plants (Cobo-Díaz *et al*. 2019, Zhou *et al*. 2016). Beyond the description of compositions and diversities of microbial communities, metabarcoding data can also be applied to predict the functionality of microbial communities (Louca *et al*. 2016; Nguyen *et al*. 2016) and examine network interactions (Vacher *et al*. 2016). The accurate description of the field microbiota, using such state-of-the-art technologies in combination with co-occurrence networks, may thus contribute to better understand the mechanisms underlying *Fusarium* spp./microbiota interactions, which has never been undertaken so far. Such knowledge may thus provide chances to develop innovative biocontrol approaches against FHB, which performance have been disappointing so far.

Indeed, it is now agreed that the soil- or plant-associated microbiota serves as a protective barrier against pathogens. While residues may represent a carrier for pathogenic organisms, including *Fusarium* spp., the removal of previous crop residues could also deprive the soil from taxa with suppressive functionalities towards such pathogens. Some candidate antagonistic organisms against *Fusarium* spp. have been isolated from maize root rhizosphere (Abiala *et al*., 2015), maize endophytes (Mosua *et al*., 2015), maize residues (Luongo *et al*., 2005; Singh *et al*., 2009) and agricultural soils (He *et al*., 2009). It was recently shown that the microbiota of maize residues were dominated by genera that contain strains previously reported as biocontrol agents, as well as plant pathogenic genera, such as *Fusarium*, *Acremonium*, *Phoma*, *Pseudomonas* and *Erwinia* (Cobo-Díaz *et al*., 2019). Therefore, a better understanding of the maize residues effects on *Fusarium* spp/ microbiota interactions may help improve crop management practices under conservation tillage systems. This could either rely on inoculative biocontrol approaches on maize residues to reduce FHB incidence or severity in following crops.

In this context, the objectives of the present work were to: i) study the dynamics of *Fusarium* and microbial communities and their interactions in soil and maize residues on agricultural fields under wheat/maize rotation; ii) determine the influence of maize residues and *F. graminearum* inoculation on soil microbial communities mesocosm conditions; iii) identify putative taxa of interest significant correlated to *Fusarium* spp. in co-occurrence networks analysis.

To address these objectives, soil and maize residue samples were collected from 8 fields in Brittany, France at 2 time-points: in November 2016 just after maize harvest and again, five months later in April 2017. Metabarcoding sequencing of *16S rRNA* gene, internal transcribed spacer (ITS) and *EF1*α gene were then used to determine the bacterial, fungal and *Fusarium* communities on those samples. In addition, our microbial community profiles were also compared to an additional time-point in April 2015 which was included in a previous study from our laboratory (Legrand *et al*., 2018). The influence of maize residues on soil microbial communities were confirmed under mesocosm conditions with soil samples inoculated or not with *Fg*.

## MATERIALS AND METHODS

### Soil and maize stalk sampling

Soils were selected from an initial amount of 31 agronomics fields sampled on April 2015 in Brittany, France (28)(Legrand et al, 2019), and a total of 8 soils were sampled in 2 additional dates, in November 2016 and April 2017. Fields localization and crop practices are described in Table 1. Fields were under maize/wheat rotation (in winter/spring rotation) for at least the last 4 years, except P16 and P23 that had no cultivation yet and onion, respectively, in sampling of April 2017. Wheat crop soils were taken one month before flowering and maize crop soils within 3 days after harvest. In each field, 15 randomly points were selected, and the first 5 cm of soil (with a hand auger of 6 cm Ø); and, for November 2016 sampling, the above-ground part of one maize stalks with nodal region and leaves were randomly sampled in each point, at the same date of soil sampling. For soil P23, where not maize residues canopy was leave on soil, some maize residues were also taken from those not collected during harvest (Fig S1). Soil samples were stored at 4°C until were sieved with a 2 mm Ø mesh before DNA extraction, that was done within 24 h after sampling. Stalks were stored at 4°C until DNA extraction, made within 1 week.

**Table 1.** Characteristics of sampled fields.

| **Code** | **Location (City)** | **GPS Coordenates** | **April 2015** | **November 2016** | **April 2017** | **Tillage** | **Fertilizers** |
| --- | --- | --- | --- | --- | --- | --- | --- |
| P06 | Gouesnou | 48.447893, -4.435354 | Wheat | Maize | Wheat | Conventional | Chemical + Manure |
| P08 | Plouvenez Lochrist | 48.598551, -4.240471 | Wheat | Maize | Wheat | Conventional | Chemical + Manure |
| P09 | Porspoder | 48.451933, -4.628142 | Wheat | Maize | Wheat | Conventional | Chemical + Manure |
| P11 | Loudéac | 48.218545, -2.806619 | Wheat | Maize | Wheat | Minimum | Chemical + Manure |
| P16 | Bannalec | 47.932287, -3.712028 | Wheat | Maize | Nothing | Conventional | Unknown |
| P20 | Plabennec | 48.500746, -4.440550 | Wheat | Maize | Wheat | Conventional | Chemical + Manure |
| P21 | Kergoz Bannalec | 47.958804, -3.705690 | Wheat | Maize | Wheat | Conventional | Chemical |
| P23 | Plouider | 48.606743, -4.281656 | Wheat | Maize * | Onion | Conventional | Chemical + Manure |

\* Maize used for silage

### Mesocosm experiment

Soils and maize samples from fields P08, P09, P11, P16, P20 and P23 were employed on mesocosm experiment. Maize residues stems were cut in pieces of around 2 cm and then crushed in a blender machine, and 5 g of maize residues were added per 100 g of soil in “Maize” treatments. The maize residues added in each soil were those picked from the same field. A *F. graminearum* inoculum, prepared according to Legrand *et al*. (2018), was added to “*Fusarium*” treatments at a proportion of 2 g of maize infected kernel per 100 g of soil. A total of 4 treatments were tested for the 6 selected fields: control soil (CS), control soil with maize residues (CM), soil with *F. graminearum* (FS) and soil with maize residues and *F. graminearum* (FM). The experiments were made within a month after sampling soils, using three replications (blocks of 4 x 4 x 6 cm) per treatment and soil, with a total of 72 blocks (4 treatments x 6 soils x 3 replicates) with 20 g of soil, or soil plus maize residues, per block. Pots were incubated in controlled conditions (day/night cycle: 16/8h, 22/18°C and 80% relative humidity) and watered each two days with sterile distilled water.

### DNA extraction

DNA was extracted from 200 mg of crushed maize stalks and leaves using FastDNA^®^ SPIN kit (MP Biomedicals, Santa Ana, CA) following the manufacturer’s instructions. For soil samples, DNA was extracted from 1 g of soil using NucleoSpin® Kit for Soil (Machery-Nagel, Dueren, De) according to the manufacturer’s instructions. Quality and concentration of purified DNA were determined using a UV spectrophotometer (NanoDrop1000, Thermo Scientific, USA).

Aliquots from environmental and mesocosm DNA extracted samples were diluted in at least 10 ng/μl before sending to the sequencing company for metabarcoding approaches.

### PCR amplification and sequencing

Soil DNA extracted at day 15 for the 4 treatments (and three replicates) in mesocosm experiment for soils P08, P09, P11, P16, P20 and P23 (72 mesocosm samples); 72 field soil samples and 24 maize samples were used for amplicon sequencing by Illumina Miseq PE300. PCR amplification, Miseq libraries preparation and sequencing was performed at the McGill University and Génome Québec Innovation Centre, Montréal, Canada. Primers 341F (5’-CCTACGGGNGGCWGCAG-3’) and 805R (5’-GACTACHVGGGTATCTAATCC-3’) (Herlemann *et al*., 2011) were used to amplify the variable regions V3 and V4 of 16S rRNA gene; primers ITS1F (5’-CTTGGTCATTTAGAGGAAGTAA-3’) and ITS4 (5’-TCCTCCGCTTATTGATATGC-3’) (White *et al*., 1990) to amplify the internal transcribed spacer; and primers Fa_150 (5’-CCGGTCACTTGATCTACCAG-3’) and Ra-2 (5’- ATGACGGTGACATAGTAGCG-3’) (Cobo-Díaz et al, 2019) to amplify the tubulin elongation factor (*ef1*α) gene of *Fusarium* species.

### Read filtering

Sequencing data were processed using QIIME (Quantitative Insights Into Microbial Ecology, version 1.9.1) (Caporaso *et al*., 2010). For 16S rDNA V3-V4 amplicons, the forward (R1) and reverse (R2) paired-end sequences were joined using *multiple_join_paired_ends.py*, followed by *multiple_split_libraries_fastq.py* for demultiplexing. Chimera sequences were removed using UCHIME algorithm (Edgar *et al*., 2011) implemented in vsearch v1.1.3 (https://github.com/torognes/vsearch) against the ChimeraSlayer database (Haas *et al*., 2011). Pick open strategy was used to cluster the sequences into Operational Taxonomic Units (OTUs) at 97% similarity cut-off using *pick_open_reference_otus.py*. The taxonomic assignment was performed using UCLUST algorithm (Edgar, 2010) against GreenGenes v13_8 database preclustered at 97% similarity cutoff (McDonald *et al*., 2012). Chloroplast, mitochondria and “No assigned” OTUs were discarded for further analysis.

The R1 and R2 paired-end sequencing reads of ITS amplicons were processed independently using *multiple_split_libraries_fastq.py*. ITS1 and ITS2 regions were first extracted separately from forward and reverse fasta files respectively, using ITSx v1.0.11 (Bengtsson-Palme *et al*., 2013) before being concatenated in a new file. A chimera filtering was made on concatenated file using the UCHIME algorithm (Edgar *et al*., 2011) with VSEARCH v1.1.3 (https://github.com/torognes/vsearch) and a modified version of the UNITE/INSDC representative/reference sequences version 7.2 (UNITE Community 2017) as reference database. The modification consisted in extracting ITS1 and ITS2 regions by ITSx software and concatenated them in the modified version of the database. The ITS1- ITS2 concatenated file of non-chimeric sequences was used for OTU picking running the QIIME script *pick_open_reference_otus.py*, with BLAST (Altschul *et al*., 1990) as taxonomic assignment method and a modified version of UNITE plus INSD non-redundant ITS database version 7.1 (Kõljalg et al 2013). Again, the modified version consisted in concatenating ITS1 and ITS2 regions after extracting them using ITSx software. Those OTUs assigned to genus *Didymella* were checked manually, by BLAST on the web service (https://blast.ncbi.nlm.nih.gov/Blast.cgi) versus nt database to improve the taxonomic assignation, and in some cases were reassigned to genus *Epicoccum,* which was not presented in the last versions of UNITE database. Only OTUs assigned to kingdom Fungi were used for further analysis. The taxonomy for fungi known to have both sexual and asexual stages was replaced by accepted names according to Chen *et al*. (2018).

The library DADA2 (Callahan *et al*., 2016) was used in R version 3.5.0 (r Development Core Team, 2017) for *ef1*α sequences filtering. Forward and reverse read pairs were trimmed and filtered, with forward reads truncated at 270 nt and reverse reads at 210 nt, no ambiguous bases allowed, and each read required to have less than two expected errors based on their quality scores. Amplicon Sequence Variants (ASVs) were independently inferred from the forward and reverse of each sample using the run-specific error rates, and then read pairs were merged requiring at least 15 bp overlap. The ASV sequences were grouped in OTUs by *pick_otus.py* QIIME script, using a 98% of similarity cutoff. OTUs representative sequences along with references were used for phylogenetic tree taxonomic assignation, according to Cobo-Díaz et al (2019).

### Statistical analysis

To minimize the inflation of rare OTUs in the community analysis, samples with less than 1,000 sequences and taxa with less than 0.01 percent relative abundance across all samples were removed, using the corresponding options in Calypso web tool (Zakrzewski *et al*., 2017). Total sum normalization and square root transformation (Hellinger transformation) were done for data normalization. Principal Coordinates analysis (PCoA) was computed with the normalized data using Bray-Curtis distance metric in Calypso web tool (Zakrzewski *et al*., 2017) while adonis test were done by v*egan* R-package (Oksanen). Richness and evenness indexes were calculated with normalized data and a previous samples rarefaction to the number of reads for the smallest sample. Wilcoxon-rank test or ANOVA test were used to compare taxa relative abundance on normalized data, and Core Microbiome analysis were computed at genus level using 0.70 as core relation samples in group. All the statistical analysis listed were made in Calypso web tool (Zakrzewski *et al*., 2017).

Metabolic and ecologically relevant functions were annotated by FAPROTAX (Louca *et al*. 2016) for the 16S rRNA gene OTU, and Wilcoxon-rank test were used to compare functions relative abundance on hellinger transformed data in Calypso web tool (Zakrzewski *et al*., 2017).

### Network analysis

The 200 most abundant OTUs were extracted from both bacterial and fungal otu-tables, and joined before were uploaded in the Molecular Ecological Network analysis Pipeline (MENAP) (Deng *et al*. 2012) in order to construct the corresponding Molecular Ecological Network (MEN) using two major steps. First, the pairwise similarity of OTU abundance across the samples was used to create a Pearson correlation matrix. Then, an adjacency matrix was determined by Random Matrix Theory (RMT)-based approach using a regressed Poisson distribution for the prediction of the nearest neighbor spacing distribution of eigenvalues.

Those nodes with the highest value of degree, betweenness, stress centrality and/or eigenvector centrality were extracted beside the nodes with a significant correlation value (edge) to plot the corresponding sub-network. Ruby homemade scripts were used to select the nodes and edges used for sub-network plots, and to add taxonomic information from OTU table to nodes files. Network graphs were plotted by Gephi 0.9.2 (Bastian *et al*. 2009) using Fruchterman Reingold spatialisation (Fruchterman *et al*. 1991). Nodes and network topological indices were calculated within the MENAP webtool (Deng *et al*. 2012).

### Accession numbers

Demultiplexed raw sequence data were deposited in the Sequence Read Archive (http://www.ncbi.nlm.nih.gov/sra) under BioProject accession number PRJNA497210, while 16S rRNA and ITS sequences from samples taken in 2015 were deposited under BioProject PRJNA429425, with Experiment numbers SRR6475734, SRR6475732, SRR6475727, SRR6475718, SRR6457721, SRR6475715, SRR6457742 and SRR6475744 for *16S rRNA*; and SRR6457737, SRR6457735, SRR6457728, SRR6475742, SRR6457747, SRR6457739, SRR6475721 and SRR6457719 for ITS region.

## RESULTS

### Shift in soil microbial communities along rotation

#### Fusarium communities

A total of 2,914,818 *ef1*α sequences were clustered into 31 OTUs (6 of them with abundance lower than 0.01 % of total *ef1*α sequences) assigned to *Fusarium* or *Neocosmospora* species, after filtering raw reads from 24 maize residues samples and 72 soil samples. Replicates 5SP09A and 7SP06C were removed for further analysis due to a low number of sequences obtained. Maize samples showed significant (p<0.05) higher richness (6.3 ± 2.1 OTUs) than soil samples (from 2.7 ± 1.6 to 4.1 ± 2.2), while no significant differences were found for evenness (Fig. 1a,b).

**Figure 1.**
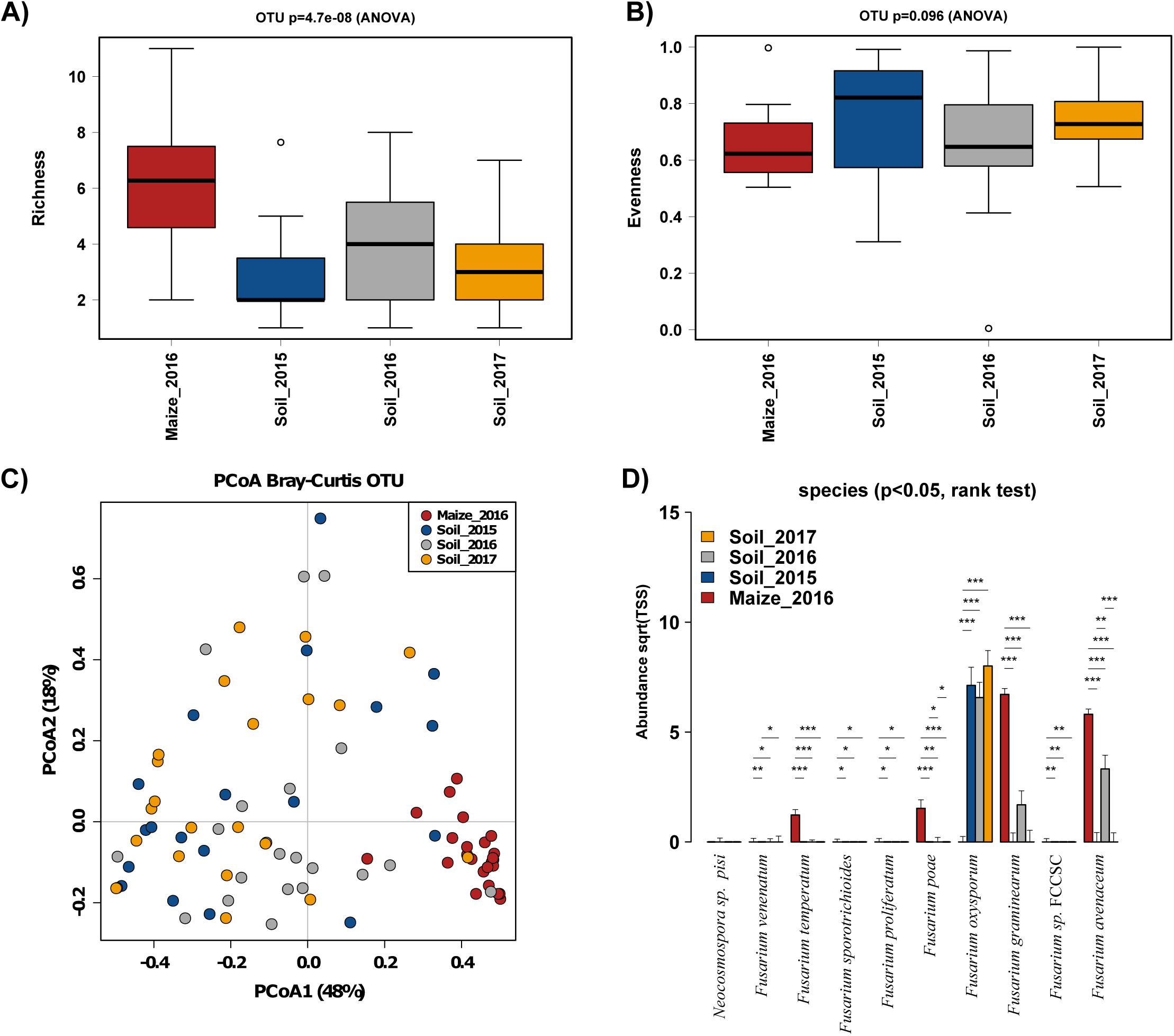
Field *Fusarium* communities. a) Richness index, b) evenness index, c) PCoA plot and d) Rank test analysis.

The *Fusarium* spp. composition of maize samples was significantly different to that of soil samples (adonis test, R^2^ = 0.29, p-value = 0.001) and there was no significant differences between soil samples (adonis test, R^2^ = 0.04, p-value = 0.084), while parcels showed significant differences for soil samples (adonis test, R^2^ = 0.32, p-value = 0.001). PCoA of *Fusarium* OTUs illustrated such differentiation among soil samples (Fig. 1c).

*F. oxysporum* was the most abundant species in soil samples, with significant higher values (rank test, p<0.05) than in maize samples while maize samples were significantly dominated by *F. graminearum* and *F. avenaceum*. In addition, *F. avenaceum* was significantly more abundant in soil samples from 2016 than the other years (Fig. 1d). Other species with significant higher abundances in maize samples than in soil samples included *F. poae*, *F. temperatum* and *Fusarium* sp. FCCSC (*Fusarium citricola* species complex), and to a lesser extent, *F. venenatum*, *F. sporotrichioides* and *F. proliferatum* (Fig. 1d).

#### Bacterial communities

A total of 2,030,793 sequences of *16S rRNA* gene were clustered into 1,753 OTUs after removing rare OTUs (relative abundance < 0.01%) from 24 maize residue and 72 soil samples. Maize samples had significant (ANOVA, p-value < 0.001) lower richness (385 ± 201 OTUs) than soil samples (from 1090 ± 57 to 1444 ± 88 OTUs) while no significant differences were observed depending on year in soil samples (Fig. 2a). Similar pattern was observed for evenness, with significant (ANOVA, p-value < 0.001) lower levels in maize samples (0.75 ± 0.05) than in soil samples (from 0.88 ± 0.03 to 0.91 ± 0.01) (Fig. 2b).

**Figure 2.**
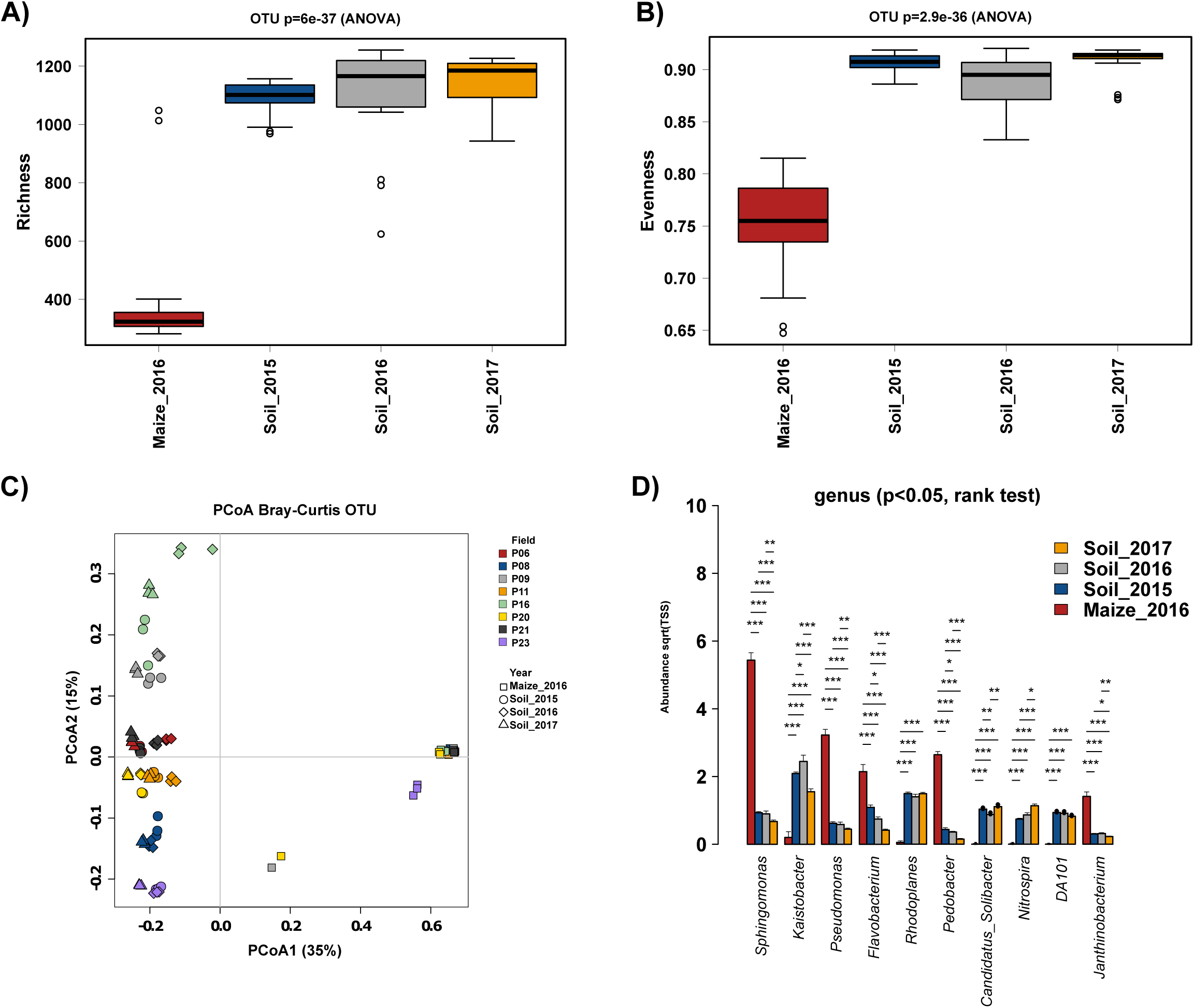
Field bacterial communities. a) Richness index, b) evenness index, c) PCoA plot and d) Rank test analysis.

Soil samples were mainly grouped by field (adonis test, R^2^ = 0.38, p-value = 0.001) in PCoA analysis, and were significant differentiated from maize samples (adonis test, R^2^ = 0.52, p-value = 0.001) (Fig. 2c).

The *Sphingomonas*, *Pseudomonas*, *Flavobacterium*, *Pedobacter*, and *Janthinobacterium* genera were significantly more abundant (rank test, p<0.05) in maize samples than in soil samples (Fig. 2d), but no increase of these genera was found on soil samples from 2016 compared to other years. The *Kaistobacter*, *Rhodoplanes*, *Nitrospira* and *DA101* genera were significantly more abundant in soil samples than in maize samples (rank test, p<0.05) (Fig. 2d).

At functional level, estimated by FAPROTAX, there were significant differences between maize and soil samples (adonis test, R^2^=0.76, p=0.001) and also within soil samples (adonis test, R^2^=0.18, p=0.001) (Fig. 3a). Some functional groups were highly represented in maize samples, including those related to chemoheterotrophy, plant pathogen and fermentation; while soil samples were enriched in functions related to nitrogen cycle and phototrophy, among others (Fig. 3b). Similar pattern was observed when comparing soil samples from 2016 to the 2 other years, which had significant higher values for chemoheterotrophy and plant pathogen in 2016, and lower values for functions related to nitrogen cycle and phototrophy (Fig. 3b).

**Figure 3.**
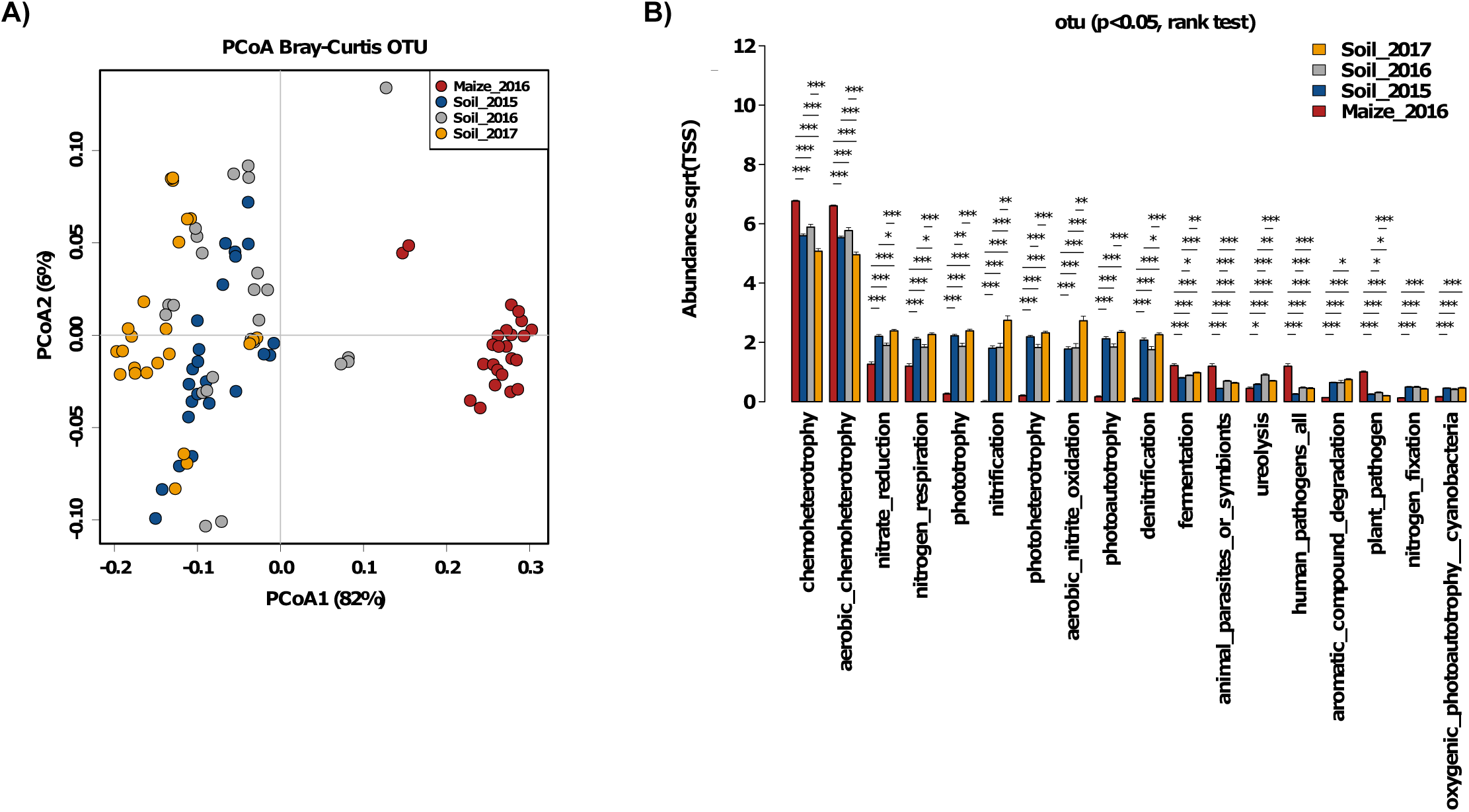
Field bacterial functionality. a) PCoA and b) Rank test analysis using the functional groups obtained by FAPROTAX pipeline.

#### Fungal communities

A total of 2,121,490 sequences of ITS were clustered into 440 OTUs after removing rare OTUs (relative abundance < 0.01%) from 24 maize residues and 72 soil samples. There was a significant increase in soil richness throughout years, with values of 69 ± 29, 113 ± 22 and 179 ± 16 OTUs for 2015, 2016 and 2017, respectively, while maize had lower richness (88 ± 15 OTUs) than soil from 2016 (Fig. 4a). Evenness values were also significantly lower in maize samples (0.40 ± 0.18) compared to soil samples (from 0.65 ± 0.08 to 0.72 ± 0.06) (Fig. 4b).

**Figure 4.**
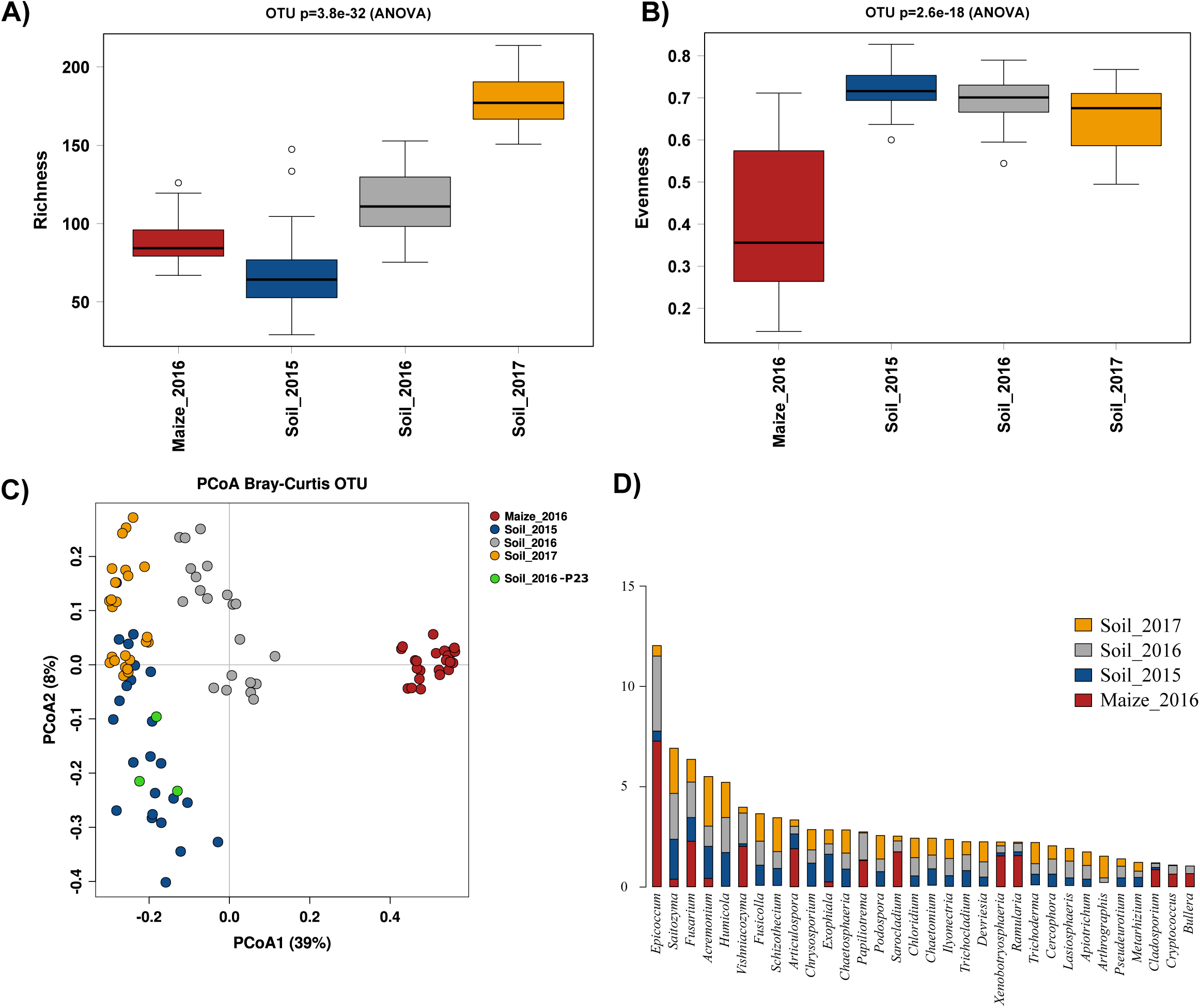
Field fungal communities. a) Richness index, b) evenness index, c) PCoA plot, d) Rank test analysis, and e) core microbiome analysis.

There was a significant effect of field (adonis test, R^2^ = 0.38, p-value = 0.001) and year (adonis test, R^2^ = 0.20 p-value = 0.001) in the composition of soil fungal communities, while soil samples were significantly diferentiate from maize residues samples (adonis test, R^2^ = 0.41, p-value = 0.001) (Fig. 4c). PCoA, at OTU level, showed that soil samples from 2016 were clustered away from the other 2 years, except for field P23 (Fig. 4c). Interestingly, this field had no maize residues left on the surface during 2016 harvest.

The *Epicoccum*, *Fusarium*, *Vishniacozyma*, *Articulospora*, *Papiliotrema*, *Sarocladium*, *Xenobotryosphaeria*, *Ramularia*, *Cladosporium*, *Cryptococcus* and *Bullera* genera were significantly more abundant (rank test, p<0.05) in maize samples than in soil samples, and, except for *Articulospora*, they were also significantly more abundant (rank test, p<0.05) in soil samples from 2016 than the other two years (Fig. 4d). Soil samples were dominated by *Saitozyma*, *Acremonium*, *Humicola*, *Fusicolla*, *Schizothecium*, *Chrysosporium* and *Exophiala* genera, with significant higher abundance in soil samples than in maize samples (Fig. 4d).

### Molecular Ecological Network analysis in field samples

Four correlation-based networks of bacterial OTUs were constructed by soil sample per year and maize residue sample. Co-occurrence network of soils from 2016 (soil_2016) showed lower connectivity (less links, average degree and connectedness) than other networks (Table 2). The lowest value for centralization of degree found in soil_2016 indicates more similarity in connectivity values within the nodes of this network, compared to others. Similar observation was found for centralization of stress centrality, which was lower in soil_2015 and soil_2016, meaning that similar values of stress centrality for the nodes within this networks (Table 2). This differences were clearly observed in the sub-network constructed with the nodes with highest value for each topological indexes (degree, stress centrality, betweenness and eigenvector centrality) and the linked nodes, where a higher proportion of nodes belonging to phylum *Gemmatimonadetes* was found for maize network, and *Actinobacteria* and *Acidobacteria* for soil networks (Fig. S2). Moreover, nodes in soil_2016 network presented a decrease in degree value (number of correlations), while some nodes presented higher eigenvalue centrality values compared to other networks (Fig. 5a). Similar observations were found for bacteria-fungi networks, with soil_2016 as the network whose nodes presented lower degree values, followed by maize_2016 and soil_2015 (Fig. 5c). This nodes with highest eigenvalue centrality were mainly *Actinobacteria* belong to *Gailellaceae* family, and no fungal node was found between them (Table S1). Some OTUs negatively correlated with nodes of *Fusarium* spp. were found, including 2 nodes in soil_2016 network (both belonging to Gammaproteobacteria), 4 nodes in soil_2017 network (belonging to Acidobacteria and Ascomycota) and 4 nodes in maize network (including one belonging to Bacteroidetes, two to Ascomycota and one to Gammaproteobacteria) (Table 3). Moreover, up to 7 nodes were positively correlated to *Fusarium* nodes in the maize network, among which 3 belonged to *Flavobacteriu*m, two to *Vishniacozyma*, one to *Sarocladium* and the other to class Sordariomycetes (Table 3).

**Figure 5.**
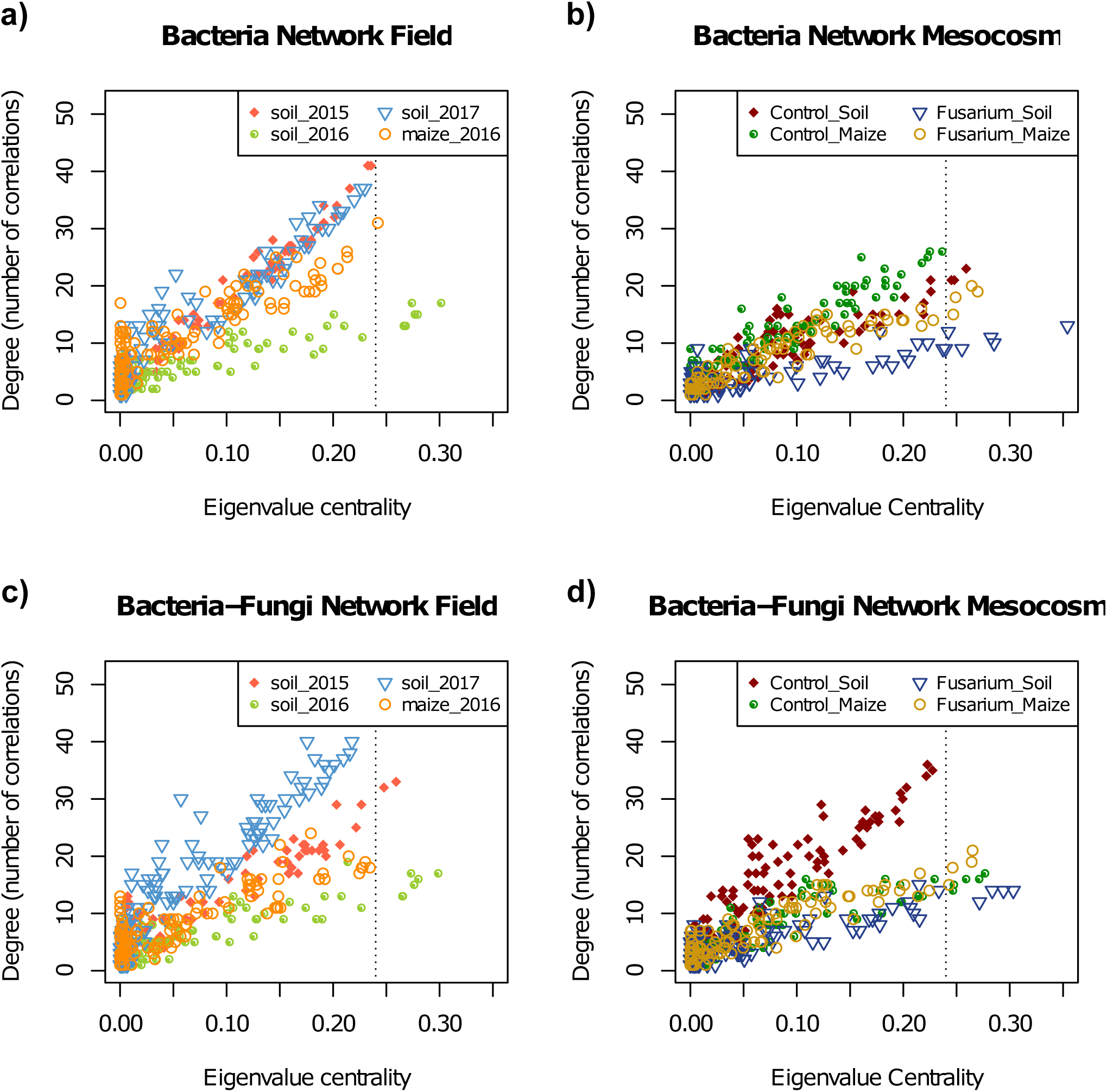
Co-occurence networks. Degree-Eigenvalue Centrality plot for nodes in a) field bacteria networks, b) mesocosm bacteria networks, c) field bacteria-fungi networks and d) bacteria-fungi networks. Vertical line indicate Eigenvalue centrality equal to 0.24.

**Table 2.** Topological properties of field bacteria networks.

| **Network Indexes** | **2015 (0.690)** | **2016 (0.820)** | **2017 (0.760)** | **maize (0.710)** |
| --- | --- | --- | --- | --- |
| Total nodes | 110 | 127 | 108 | 145 |
| Total links | 661 | 304 | 698 | 763 |
| R square of power-law | 0.496 | 0.909 | 0.39 | 0.305 |
| Average degree (avgK) | 12.018 | 4.787 | 12.926 | 10.524 |
| Average clustering coefficient (avgCC) | 0.449 | 0.346 | 0.473 | 0.598 |
| Average path distance (GD) | 2.699 | 3.747 | 3.044 | 3.935 |
| Geodesic efficiency (E) | 0.465 | 0.34 | 0.421 | 0.333 |
| Harmonic geodesic distance (HD) | 2.149 | 2.939 | 2.373 | 3.006 |
| Maximal degree | 41 | 17 | 37 | 31 |
| Centralization of degree (CD) | 0.271 | 0.098 | 0.229 | 0.144 |
| Maximal betweenness | 453.174 | 1312.223 | 1300.332 | 1717.985 |
| Centralization of betweenness (CB) | 0.066 | 0.156 | 0.213 | 0.147 |
| Maximal stress centrality | 3426 | 4244 | 12784 | 39741 |
| Centralization of stress centrality (CS) | 0.48 | 0.487 | 2.038 | 3.5 |
| Maximal eigenvector centrality | 0.236 | 0.301 | 0.23 | 0.242 |
| Centralization of eigenvector centrality (CE) | 0.174 | 0.257 | 0.168 | 0.193 |
| Density (D) | 0.11 | 0.038 | 0.121 | 0.073 |
| Transitivity (Trans) | 0.568 | 0.416 | 0.604 | 0.569 |
| Connectedness (Con) | 0.748 | 0.56 | 0.963 | 1 |
| Efficiency | 0.863 | 0.945 | 0.883 | 0.933 |

**Table 3.**
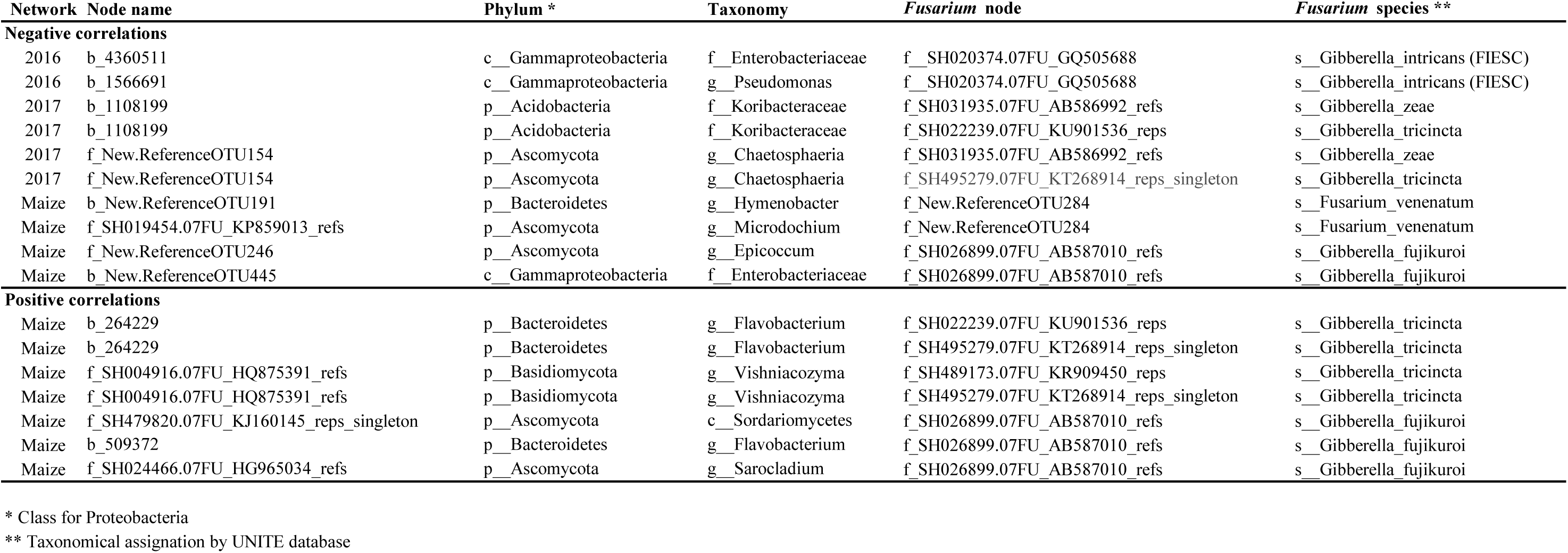
Nodes in field networks with significant co-occurrence with *Fusarium* spp. nodes

### Influence of maize residues and *F. graminearum* inoculation in soil microbial communities under mesocosm conditions

#### Bacterial communities

A total of 1,689,356 sequences of *16S rRNA* gene were clustered into 1,822 OTUs after removing rare OTUs (relative abundance < 0.01%) from 72 mesocosm soil samples. Excluding samples from field P16, which presented lowest values for alpha diversity indices, *Fg* inoculation or amendment with maize residues induced higher richness than in control samples (1289 ± 61 OTUs in SC compared to 1367 ± 54 OTUs1341 ± 55 OTUs, in MC and SF respectively)(Fig. S3a). When comparing samples amended with maize residues, Fg inoculation induced significant lower evenness values, (0.85 ± 0.06 versus 0.90 ± 0.01, in MF and MC, respectively)(Fig. S3b).

In terms of bacterial compositional structure, differences due to treatment (adonis test, R^2^ = 0.09, p- value = 0.007) were much less than the variation between fields (adonis test, R^2^ = 0.70, p-value = 0.001), as was shown in PCoA plot (Fig. 6a).

**Figure 6.**
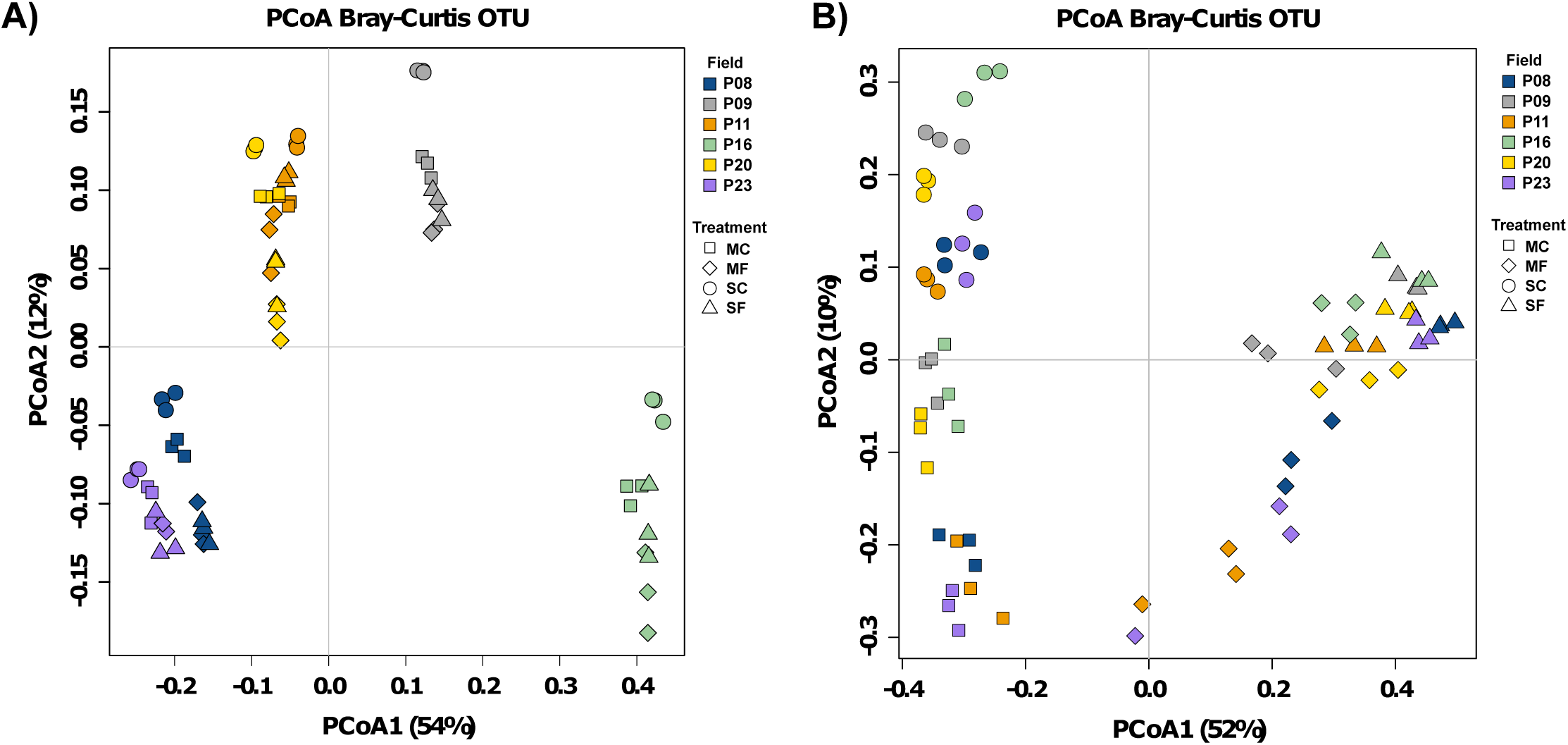
Mesocosm betadiversity analysis. PCoA plot of a) bacterial and b) fungal communities obtained in mesocosm experiment at d15.

The addition of maize residues induced a significant increase in *Sphingomonas*, *Opitutus*, *Chthoniobacter* and *Fimbriimonas* relative abundance and a decrease in *Methylibium*, *Nitrospira* and *Terracoccus* whether or not soils were inoculated with Fg (Fig. S3c,d). In non-inoculated soil, the addition of maize induced a significant higher levels of other genera such as *Cellvibrio*, *Devosia*, *Phenylobacterium*, *Luteolibacter*, *Caulobacter*, *Luteibacter* and *Prosthecobacter* and lower levels for *Kaistobacter*, *Pseudomonas* and *Nitrospira* genera (Fig. S3d). Whether or not soil were amended with maize residues, *Fg* inoculation induced an increase in relative abundance for *Sphingomonas*, *Burkholderia*, *Pseudomonas*, *Rhodanobacter*, *Cellvibrio*, *Opitutus* and *Rhizobium*, while *Kaistobacter* and *Rhodoplanes* were decreased (Fig. S3e).

At functional level, estimated by FAPROTAX, the addition of maize residues increased the relative abundance of OTUs assigned to nitrogen fixation, fermentation and cellulolysis, among other functions (Fig. S4a). Fg inoculation induced an increase in nitrogen fixation and plant pathogen functions, among others, and a decrease in others functions associated to nitrogen metabolism, such as nitrogen respiration in soil treatments (Fig. S4b) and nitrogen respiration, nitrate reduction and denitrification in maize residues treatments (Fig. S4c).

#### Fungal communities

A total of 2,432,998 sequences of ITS were clustered into 264 OTUs after removing rare OTUs (relative abundance < 0.01%) from 72 mesocosm soil samples. Richness was significantly higher in MC treatment, with 40 ± 16 OTUs in SF, 144 ± 27 in SC, 79 ± 20 in MF and 136 ± 21 in MC (Fig. S5a). No significant differences were found for evenness values, which ranged from 0.60 ± 0.11 to 0.71 ± 0.06 (Fig. S5b).

PCoA showed a significant clustering of samples by treatment (adonis test, R^2^ = 0.62, p-value = 0.001), while slight differences were found by field (adonis test, R^2^ = 0.12, p-value = 0.04) (Fig. 6b).

A significant decrease in *Fusarium* relative abundance was found in MF treatment compared to SF, with significant higher relative abundance of *Epicoccum*, *Podospora*, *Lasiosphaeris* and *Sarocladium* genera, among others (rank test, p<0.05), in MF compared to SF (Fig. S4c). In non-inoculated soils, the addition of maize (MC treatment) induced an increase in the *Podospora*, *Sarocladium*, *Ramularia*, *Cryptococcu*s, *Phaeoacremonium* and *Apodus* genera (rank test, p<0.05) and a decrease in the *Acremonium*, *Fusicolla*, *Humicola* and *Ilyonectria* genera, among others, compared to SC treatment (Fig. S4d).

### Molecular Ecological Network analysis in mesocosm experiment

After removing samples for soil P16, which presented an important increase of *Rhodanobacter* genus for all the treatments (data not shown), co-occurrence network analysis were constructed for both bacterial and fungal data pooled together, and with bacterial data only. In bacteria networks, CM treatment showed the highest values for total links and average degree, with a progressive decrease for CS, FM and FS networks for both parameters (Table 4). Moreover, FS treatment presented the lowest values for average clustering coefficient, centralization of degree and density, while had the highest values for average path distance, harmonic geodesic distance, centralization betweenness and centralization of eigenvector centrality (Table 4). Lower values for the degree eigenvalue centrality proportion was observed in nodes from FS in bacteria networks (Fig. 5b) and in nodes from all treatments except CS for bacteria-fungi network (Fig. 5d). Nodes with highest eigenvalue centrality were mainly assigned to Kaistobacter genus and/or Proteobacteria phylum, and no fungal node was detected within them (Table S1).

**Table 4.** Topological properties of mesocosm bacteria networks.

| **Network Indexes** | **CM (0.860)** | **CS (0.880)** | **FM (0.860)** | **FS (0.900)** |
| --- | --- | --- | --- | --- |
| Total nodes | 131 | 111 | 112 | 93 |
| Total links | 557 | 410 | 351 | 196 |
| R square of power-law | 0.642 | 0.598 | 0.695 | 0.758 |
| Average degree (avgK) | 8.504 | 7.387 | 6.268 | 4.215 |
| Average clustering coefficient (avgCC) | 0.447 | 0.458 | 0.425 | 0.346 |
| Average path distance (GD) | 3.664 | 3.147 | 3.596 | 4.234 |
| Geodesic efficiency (E) | 0.364 | 0.394 | 0.363 | 0.304 |
| Harmonic geodesic distance (HD) | 2.748 | 2.539 | 2.758 | 3.286 |
| Maximal degree | 26 | 23 | 20 | 13 |
| Centralization of degree (CD) | 0.137 | 0.145 | 0.126 | 0.098 |
| Maximal betweenness | 1396.241 | 716.186 | 743.481 | 1320.59 |
| Centralization of betweenness (CB) | 0.152 | 0.106 | 0.106 | 0.289 |
| Maximal stress centrality | 20169 | 8982 | 3948 | 4020 |
| Centralization of stress centrality (CS) | 2.201 | 1.344 | 0.546 | 0.878 |
| Maximal eigenvector centrality | 0.237 | 0.259 | 0.27 | 0.354 |
| Centralization of eigenvector centrality (CE) | 0.181 | 0.194 | 0.211 | 0.298 |
| Density (D) | 0.065 | 0.067 | 0.056 | 0.046 |
| Transitivity (Trans) | 0.527 | 0.442 | 0.439 | 0.409 |
| Connectedness (Con) | 0.759 | 0.752 | 0.721 | 0.836 |
| Efficiency | 0.923 | 0.921 | 0.932 | 0.957 |

Finally, fungi-fungi intra-kingdom and fungi-bacteria inter-kingdom connections were higher in mesoscosm treatments, except for FS treatment for f-f conections, compared to environmental networks,. (Table 5). Moreover, presence of maize residues decrease the levels of the 3 kind of connections (bacteria-bacteria, fungi-fungi, and fungi-bacteria) for soil environmental and mesocosm networks, except for ff connection in mesocosm (Table 5).

**Table 5.** Analysis of the proportion of intra-kingdom interactions in the co-occurrence networks.

| Sample | Number of nodes |  | Number of interactions |  |  | Theoretical Max. interactions |  |  | % of interactions <sup>4</sup> |  |  |
| --- | --- | --- | --- | --- | --- | --- | --- | --- | --- | --- | --- |
|  | bact | fung | bb | ff | fb | bb <sup>1</sup> | ff <sup>2</sup> | fb <sup>3</sup> | bb | ff | fb |
| 2015 | 95 | 112 | 440 | 3 | 74 | 4465 | 6216 | 10640 | 9.85 | 0.05 | 0.70 |
| 2016 | 132 | 160 | 304 | 4 | 50 | 8646 | 12720 | 21120 | 3.52 | 0.03 | 0.24 |
| 2017 | 112 | 181 | 698 | 58 | 269 | 6216 | 16290 | 20272 | 11.23 | 0.36 | 1.33 |
| Maize | 138 | 166 | 500 | 27 | 68 | 9453 | 13695 | 22908 | 5.29 | 0.20 | 0.30 |
| FM | 114 | 22 | 351 | 10 | 48 | 6441 | 231 | 2508 | 5.45 | 4.33 | 1.91 |
| FS | 112 | 9 | 267 | 0 | 20 | 6216 | 36 | 1008 | 4.30 | 0.00 | 1.98 |
| CS | 130 | 44 | 680 | 17 | 137 | 8385 | 946 | 5720 | 8.11 | 1.80 | 2.40 |
| CM | 113 | 40 | 317 | 23 | 49 | 6328 | 780 | 4520 | 5.01 | 2.95 | 1.08 |

<sup>1</sup> estimated by $x = n! / (2x(n-1)!)$ , where $n$ is the number of bacteria nodes
<sup>2</sup> estimated by $x = n! / (2x(n-1)!)$ , where $n$ is the number of fungi nodes
<sup>3</sup> estimated by $x = n_f \times n_b$ , where $n_f$ and $n_b$ are the number of fungi and bacteria nodes, respectively
<sup>4</sup> percentage of interactions found from the theoretical maximum interactions

## DISCUSSION

Basic but yet unanswered questions regarding the ecology of *Fusarium* spp. still remain. First, although soil and residues constitute the main FHB inoculum sources (Bateman *et al*. 2007; Fernandez *et al*. 2008), the knowledge on microbial ecology in residues and its influence on pathogen dispersion or suppression is scarce. These findings could be important for the selection of appropriate control strategies in order to determine the predominant *Fusarium* spp. that should be targeted and the field components and wheat stages during which treatments are more likely to affect pathogen populations or their impact on the plant. In the present study, *F. graminearum*, *F. avenaceum, F. poae* and *F. temperatum* were the most abundant *Fusarium* species found on maize residues, as has already been reported on maize stalks after a 6-month field exposure (Köhl et al, 2015), maize residues collected after harvest (Cobo-Diaz *et al*. 2019, Dill-Macky and Jones 2000), maize stalks and kernels (Basler 2016), wheat kernels (Xu *et al*. 2005; Karlsson *et al*. 2016, 2017; Nicolaisen *et al*. 2014) and barley kernels (Schöneberg *et al*. 2016). In contrast, *F. oxysporum* was found as the main species in soil samples, as has been found previously on wheat crops (Edel-Hermann et al, 2015; Silvestro *et al*., 2013; Leblanc et al, 2015). Moreover, an important input of *F. graminearum* and *F. avenaceum* from maize residues to soils was found, which confirmed maize residues as an important source of these *Fusarium* species associated to FHB. Furthermore, the low abundance of this species found in soils sampled before wheat flowering, after maize inter-crops and conventional tillage (except P11 field, which was under minimum tillage, and P23, with not maize residues canopy), support the idea of use conventional tillage approaches to reduce the availability of *Fusarium* spp. pathogens for following crops, as has been suggested before (Schöneberg *et al*. 2016).

Those fungal genera promoted by maize residues addition harbour maize or/and wheat pathogens species, such as *Cladosporium, Fusarium* and *Epicoccum*; and other plant pathogens, such as *Sarocladium* and *Ramularia*. Conversely, strains with antagonism effect against *F. graminearum* in wheat were found for *Epicoccum* (Luongo *et al*. 2005; Jensen *et al*., 2016)*, Cladosporium* (Luongo *et al*. 2005) and *Sarocladium* (Comby *et al*. 2017). Another genera deposited in soil from maize residues were *Vishniacozyma*, *Papiliotrema*, *Cryptococcus* and *Bullera*, which belong to order *Tremellales*.

Some strains of *Cryptococcus* were described as effective biocontrol agent (BCA) against *Fusarium* spp. in wheat (Wachowska *et al*. 2013b; Schisler *et al*. 2011) and those genera belong to *Tremellales* could be considered within this group of putative BCA against *Fusarium* spp. because they contain strains previously grouped within *Cryptococcus* genus (Liu *et al*. 2015). This field results were supported with the results obtained in mesocosm experiments, which reported also a increase in relative abundance due to maize residues of genera that harbor strains reported previously as BCA or organism associated to healthy soils in *Fusarium* spp. diseases studies, such as *Cryptococcus* (Wachowska *et al*. 2013b; Schisler *et al*. 2011), *Articulospora* (Sugahara *et al*. 2018) and *Sarocladium* (Comby *et al*. 2017). Furthermore, maize residues microbial co-occurrence networks presented some OTUs assigned to *Sarocladium*, *Epicoccum* and *Vishniacozyma* genera with significant correlation with OTUs assigned *Fusarium* spp. which could be an evidence of antagonism effect of this genera versus *Fusarium* spp. The same results, except for *Vishniacozyma*, were found for another maize residues sampling in Brittany, France (Cobo-Díaz *et al*. 2019), and their relative abundance increase in soils could have a antagonistic effect against *Fusarium* spp. *Flavobacterium,* S*phingomonas, Pseudomonas*, *Pedobacter* and *Janthinobacterium* were found as the most abundant in our maize samples, as has been reported previously for maize residues (Cobo-Díaz *et al*. 2019), maize rhizospheric soils (Yang *et al*. 2017a; García-Salamanca *et al*. 2013; Li *et al*. 2014; Correa-Galeote *et al*. 2016) or even in wheat rhizospheric soils (Yin *et al*. 2013). This bacterial genera were reported as bacterial genera associated with reduced colonization of *Fusarium* spp. in maize stalks (Köhl *et al*. 2015) and/or contains strains characterized as BCA against *Fusarium* spp. (Wachowska *et al*. 2013a,b; Ito *et al*. 2013; Chen *et al*. 2018A; De Boer *et al*. 2007; Haack *et al*. 2016). The predominance in maize residues of such genera make them a potential source of bacterial species for plant pathogen control, but no increase of their relative abundance in the corresponding soil samples were observed. Maybe it could be due to the short time lapsed between harvest and sampling, which was no more than 3 days. Furthermore, maize residues generate changes in the soil nitrogen transformations (Li *et al*. 2019) and increase soil nitrogen content (Maresma *et al*. 2018), and could generate the decrease, in relative abundance, of some bacterial functions related to nitrogen metabolism observed in our field samples.

Differences in the soil bacterial composition structure among years was lower than variation between fields, while fungal communities composition were significant different between soil samples from 2016 and the other 2 years, except for P23, whose samples from 2016 were within the 2015-2017 group. This field was the only one not amened with maize residues, although the base of the stalks were leave on crop (see Fig. S1). This maize residues influence on fungal communities observed in PCoA analysis was highlighted with the increase of relative abundance in soils after maize harvest of many fungal genera that clearly comes from those residues left on the field, while not increase was found on bacterial genera due to maize residues. Moreover, mesocosm experiment results corroborated this important influence of maize residues (and also *Fg* inoculation) on fungal communities structure and composition, while not influence was found for bacteria.

This stronger influence in fungal than bacterial communities had been observed in soil transplant experiments along 6 years (Zhao *et al*., 2019). Moreover, it have been found that microbial diversity or taxonomical information are not sensitive enough as indicator of ecosystem perturbations (Karimi *et al*. 2016) and in some cases significant changes in bacterial co-occurence patterns were found while no differences were observed in diversity indexes (He *et al*. 2017). The use of ecological networks as indicators of environmental quality had reported that higher levels of perturbation correlated to lower complexity in microbial networks (Karimi *et al*. 2016, Lupatini *et al*. 2014, Zappelini *et al*. 2015, Sauvadet *et al*. 2016), including disease incidence as perturbation factor (Yang *et al*., 2017b). In our study, although bacterial communities did not present strong differences due to maize presence, or even along years of sampling, their co-occurrence networks presented a significant decrease of connectivity and nodes degree values when maize were added to soil samples. The main factor could be changes in interaction between species due to the increase of nutrients and also the differences in fungal communities composition observed by maize residues addition. Moreover, this changes on bacterial networks was accentuated with the increase of importance for nodes belonging to *Actinobacteria* phylum, which although were not abundant in the communities, had an important role within co-occurence networks. Phylum *Actinobacteria* was found previoulsy as a key taxon on bacterial soil networks, where it could reduce the chance of soil plant pathogen invasion for tobacco bacterial wilt disease (Yang *et al*. 2017b). The decrease of connectivity values has also been observed on soil bacterial communities due to land use, where higher density of links were found in natural forest soils than pasture or field and plantations soils (Lupatini *et al*. 2014), so that maize addition could be considered as a strong perturbaction of soil microbial co-occurrence networks. Furthermore, the not clearly existence of keystone taxa (data not shown) could be advantageous to the ecosystem functionality, as the lost or decrease on any microbial taxa is not going to weaken the inter correlation network of the ecosystem (Toju *et al*. 2018).

## CONCLUSIONS

Maize residues have a stronger influence in fungal communities than bacterial communities composition, although reduction on connectivity indexes for bacterial co-occurrence networks was found due to maize addition. Bacterial communities composition were conserved along the time, with a clear differentiation due to the field instead of time of sampling. Maize residues harbour both bacterial and fungal genera previously reported as biocontrol agents against *Fusarium* spp. or diseases caused by this genera, but only an increase in relative abundance of those genera belonged to fungi were observed in both field and mesocosm soils amened with maize residues. Some OTUs belonged to those BCA genera, such as the bacteria *Flavobacterium* and the fungal *Epicoccum*, *Vishniacozyma* and *Sarocladium*, presented significant co-occurrence with *Fusarium* spp. OTUs, mainly in maize networks. Further experiments and studies has to be done to clarify their effect in *Fusarium* disease suppression in following crops, and to found which agricultural practices can increase the presence of those BCA genera in soil, and reduce of FHB events or severity.

---

**Table S1. Nodes topological characteristics and taxonomic assignment.** Only nodes with highest eigenvalue centrality (> 0.24) were indicated. “Network” column indicates to which networks belong (year number for field soil networks, “Maize” for maize network, treatment for mesocosm networks).

**Figure S1. Sampled fields in November 2016.** Photos of sampled fields a) P08, b) P09, c) P20 and d) P23. Maize residues from P23 were used for silage and not left in crop.

**Figure S2. Bacteria co-occurrence sub-network.** Nodes with highest topological characteristics (plotted in yellow) and those correlated to them were plotted. Nodes were colored by phylum and edges according to positive (green) or negative (red) correlation between nodes linked.

**Figure S3. Mesocosm bacterial communities.** a) Richness index, b) evenness index, c) Rank test analysis for *Fusarium* treatments, d) Rank test analysis for Control treatments and e) Rank test analysis compared *Fusarium* treatments versus control treatments.

**Figure S4. Mesocosm bacterial functionality.** a) Rank test analysis for control treatments, b) soil treatments and c) maize treatments, using the functional groups obtained by FAPROTAX pipeline.

**Figure S5. Field fungal communities.** a) Richness index, b) evenness index, c) Rank test analysis for *Fusarium* treatments, d) Rank test analysis for Control treatments.

