## Supplementary material for "Maize residues changes soil fungal composition and decrease soil microbial co-ocurrence networks complexity": Fig. S1

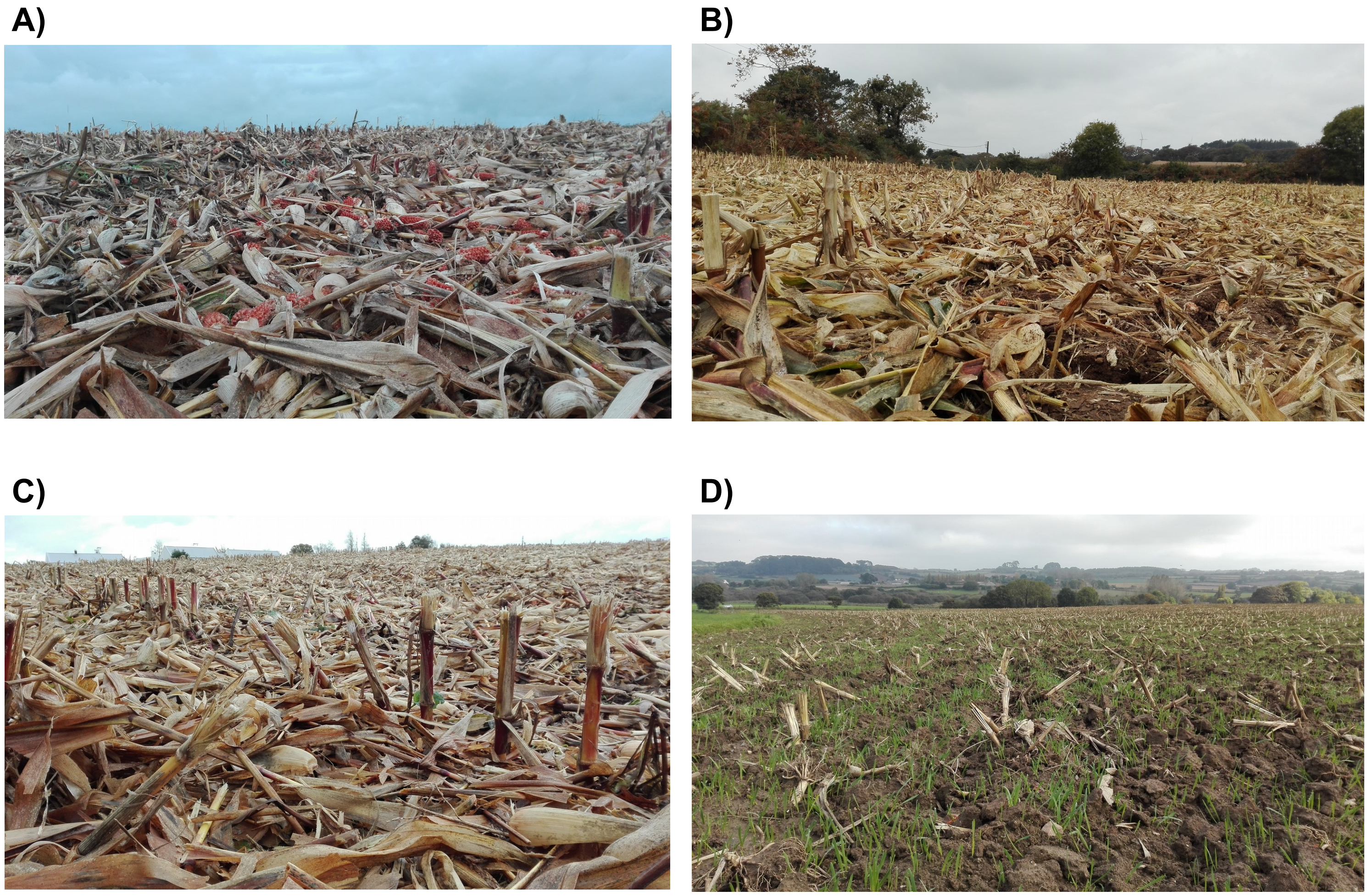


**Figure S1. Sampled fields in November 2016.** Photos of sampled fields a) P08, b) P09, c) P20 and d) P23. Maize residues from P23 were used for silage and not left in crop.
