## Supplementary material for "Maize residues changes soil fungal composition and decrease soil microbial co-ocurrence networks complexity": Fig. S2

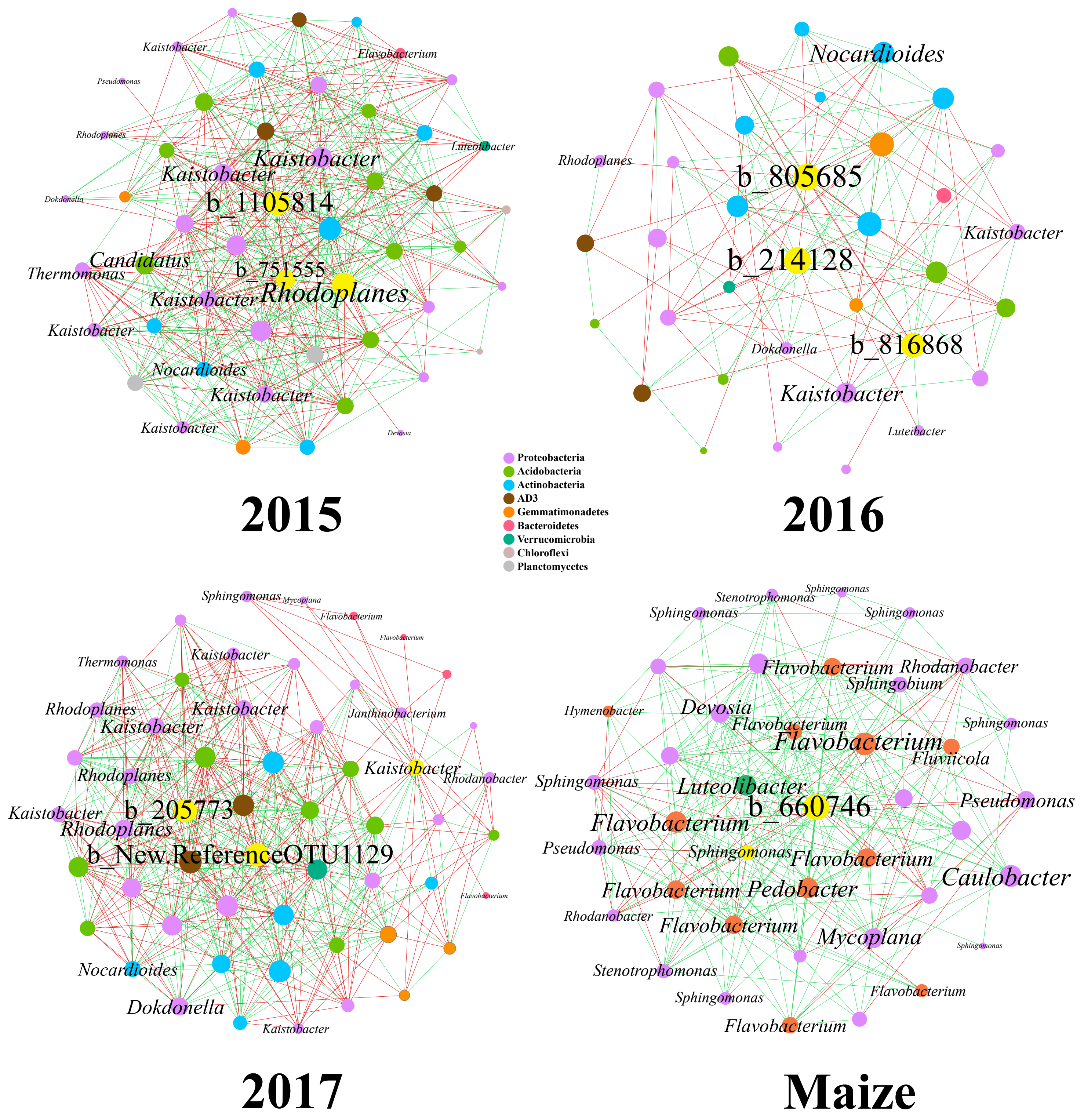


**Figure S2. Bacteria co -occurrence sub-network.** Nodes with highest topological characteristics (plotted in yellow) and those correlated to them were plotted. Nodes were colored by phylum and edges according to positive (green) or negative (red) correlation between nodes linked.
