## Supplementary material for "Maize residues changes soil fungal composition and decrease soil microbial co-ocurrence networks complexity": Fig. S3

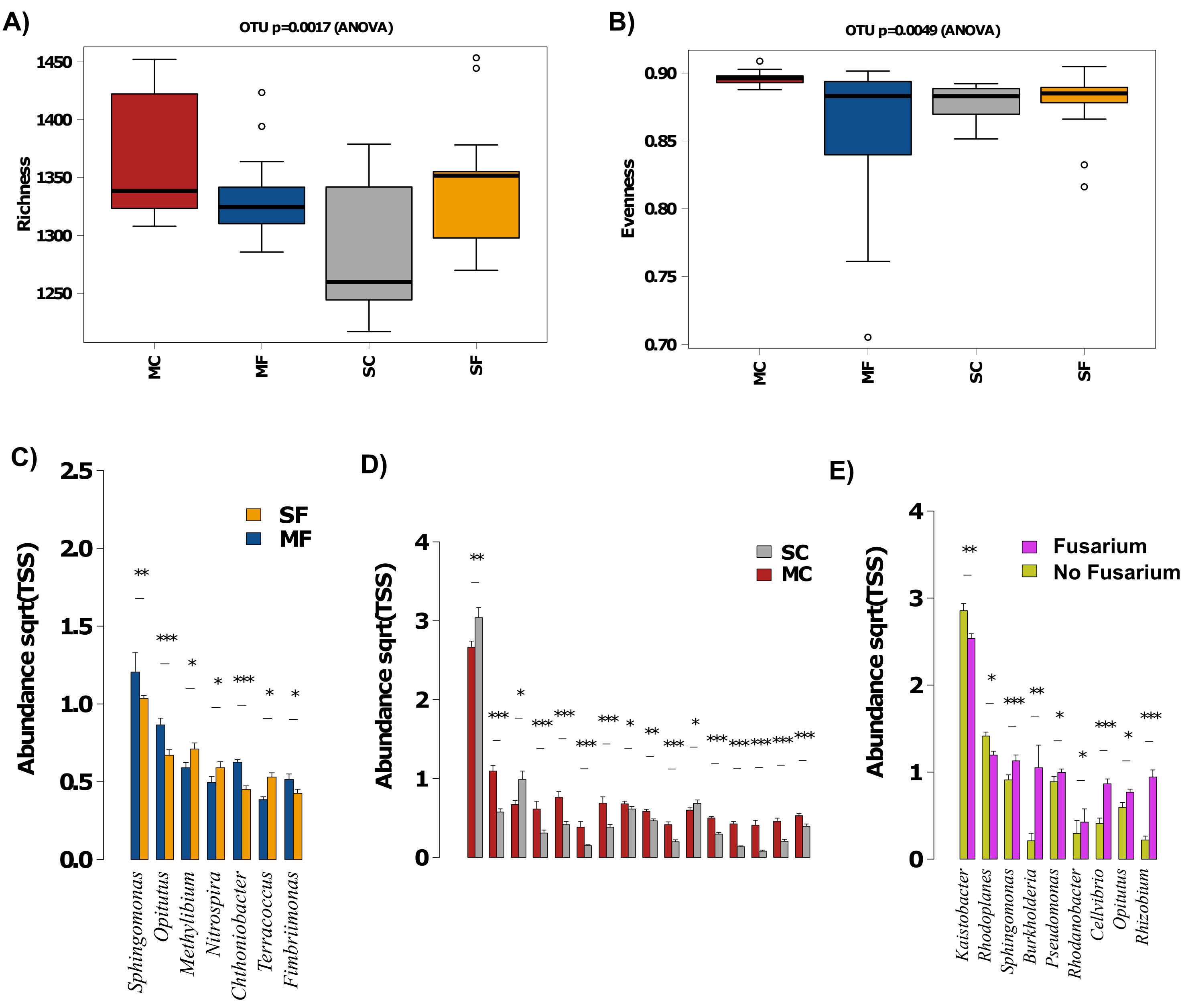


**Figure S3. Mesocosm bacterial commun ities.** a) Richness index, b) evenness index, c) Rank test analysis for *Fusarium* treatments, d) Rank test analysis for Control treatments and e) Rank test analysis compared *Fusarium* treatments versus control treatments.
