## Supplementary material for "Maize residues changes soil fungal composition and decrease soil microbial co-ocurrence networks complexity": Fig. S4

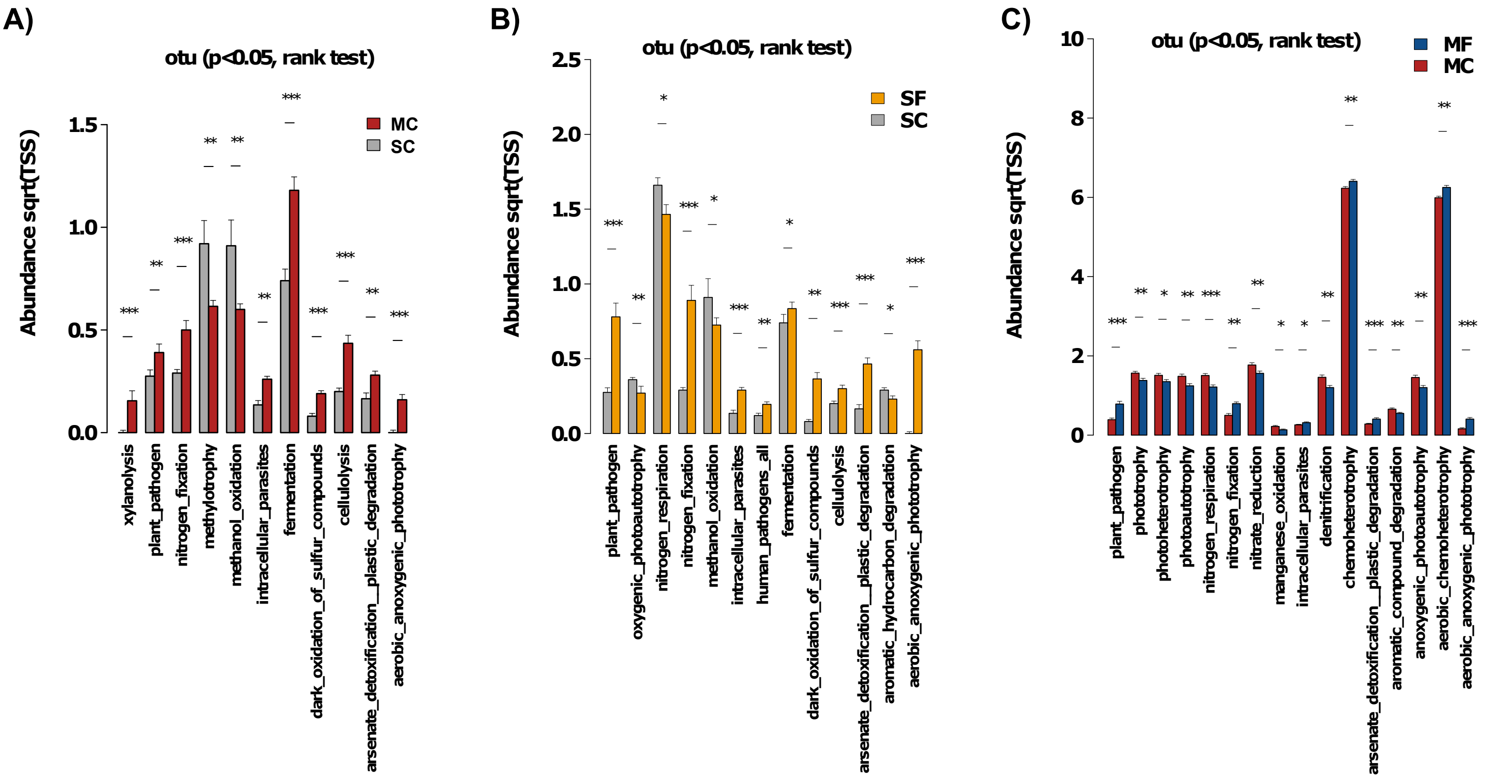


**Figure S4. Mesocosm bacterial functio nality.** a) Rank test analysis for control treatments, b) soil treatments and c) maize treatments, using the functional groups obtained by FAPROTAX pipeline.
