## Supplementary material for "Maize residues changes soil fungal composition and decrease soil microbial co-ocurrence networks complexity": Fig. S5

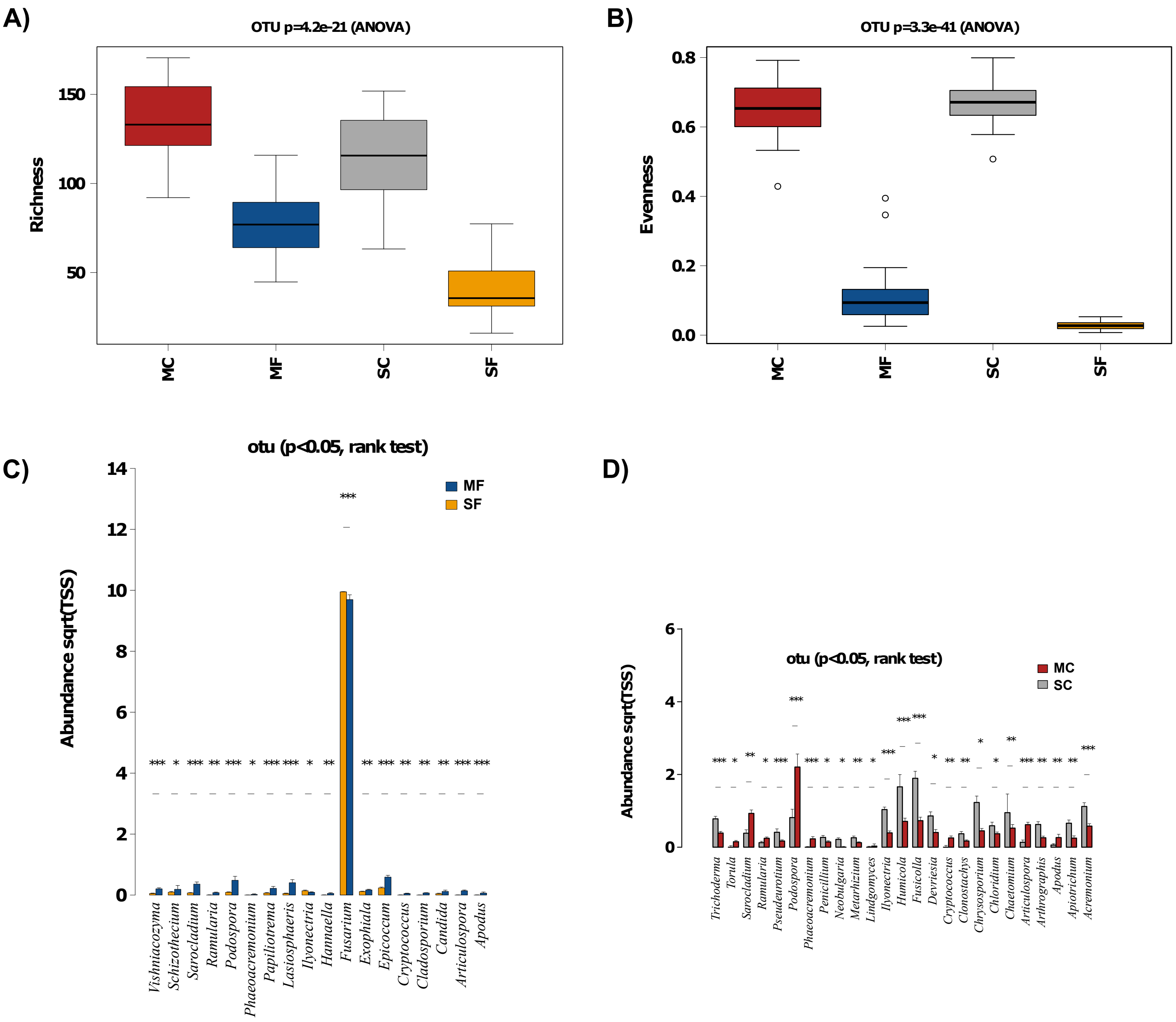


**Figure S5. Field fungal communities.** a) Richness index, b) evenness index, c) Rank test analysis for *Fusarium* treatments, d) Rank test analysis for Control treatments.
